## Supplementary Information for "A step towards measuring connectivity in the deep sea: elemental fingerprints of mollusk larval shells discriminate hydrothermal vent sites"

**Table of contents:**


**Page 2: Supplementary Information S1: Geographic classes and distributions of abundances for the entire *Shinkailepas tollmanni* larval shell dataset**

**Page 4: Supplementary Information S2: Laserogram of acquisition of composition of encapsulated larval shell**

**Page 5: Supplementary Information S3: Best classification models for *Shinkailepas tollmanni* larval shells elemental fingerprinting**

**Page 6: Supplementary Information S4: Matlab scripts and functions for randomization**

**Page 34: Supplementary Information S5: Elemental composition of the habitat water surrounding the *Ifremeria nautilei* communities sampled at hydrothermal-vent sites in the Southwest Pacific**

**Page 36: Supplementary Information S6: Boxplots for each element measured in the shells of *Shinkailepas tollmanni* encapsulated larvae**

**Page 40: Supplementary Information S7: Details on the larval shell elemental distributions in the Lau area**

**Supplementary Information S1**

**Geographic classes and distributions of abundances for the entire *Shinkailepas tollmanni* larval shell dataset**

A total of 600 individual encapsulated larval shells was sampled and analyzed from 14 hydrothermal sites, separated into areas and regions. Although part of the Lau Basin, the Mangatolo sites were separated in our study due to the large distance between this area and the three other sites from the Lau Basin. For some sites, *Shinkailepas tollmanni* capsules were collected from several spots (referred to as 'bioboxes'), not necessarily from the same chimney.

**Table S1:** Coordinates of the sampling sites.

| Area | Site | Coordinates |
|---|---|---|
| Manus | Roman Ruins | 3°43'39" S, 151°40'51" E |
| | Fenway | 3°43'42" S, 151°40'20" E |
| | Solwara 8 | 3°48'49" S, 151°40'27" E |
| | North Su | 3°47'56" S, 152°06'02" E |
| | South Su | 3°48'29" S, 152°06'17" E |
| Woodlark | La Scala | 9°47'57" S, 155°03'10" E |
| North Fiji | Phoenix | 16°57'00" S, 173°55'08" E |
| Futuna | Kulo Lasi | 14°56'28" S, 177°15'33" W |
| | Fatu Kapa | 14°44'15" S, 177°09'58" W |
| Lau | Tu'i Malila | 21°59'21" S, 176°34'06" W |
| | ABE | 20°45'47" S, 176°11'28" W |
| | Tow Cam | 20°19'05" S, 176°08'16" W |
| Mangatolo | Mangatolo North | 15°24'52" S, 174°39'12" W |
| | Mangatolo South | 15°24'57" S, 174°39'20" W |

**Table S2:** Number of individual larvae analyzed for each hydrothermal site. Some sites were sampled more than once and the number of larvae for each biobox is indicated in parentheses.

| Region | Area | Site | Number of bioboxes | Number of individual larvae |
|---|---|---|---|---|
| East | Lau | Tow Cam | 2 | **10** (2+8) |
| | | Tu'i Malila | 2 | **49** (40+9) |
| | | ABE | 2 | **73** (38+35) |
| | Mangatolo | North | 1 | **50** |
| | | South | 1 | **24** |
| | Futuna | Kulo Lasi | 1 | **46** |
| | | Fatu Kapa | 4 | **141** (53+21+34+33) |
| | North Fiji | Phoenix | 2 | **96** (48+48) |
| West | Manus | Fenway | 1 | **19** |
| | | Solwara | 2 | **44** (16+28) |
| | | South Su | 2 | **26** (2+24) |
| | | North Su | 2 | **11** (6+5) |
| | | Roman Ruins | 1 | **7** |
| | Woodlark | Scala | 1 | **4** |

Metal abundances in the 600 analyzed shells of *S. tollmanni* encapsulated larvae are presented in **Figure S1**. From these, 104 shells presented one or more measurements below the limit of detection (LOD), altogether representing 124 measurements (all elements combined) out of 7800 (1.59%). These measurements corresponded to values of As (33 shells), Cr (30 shells), Cd (30 shells), Sb (30 shells), and Cu (1 shell). We opted to keep these measurements below the LOD for comparison purposes in subsequent data analyses as these low abundances are informative, albeit not quantitative. Their values have been assigned the arbitrary value of 3.85 $10^{-8}$ µg.$g^{-1}$ to reflect a much lower value than the rest of the dataset.

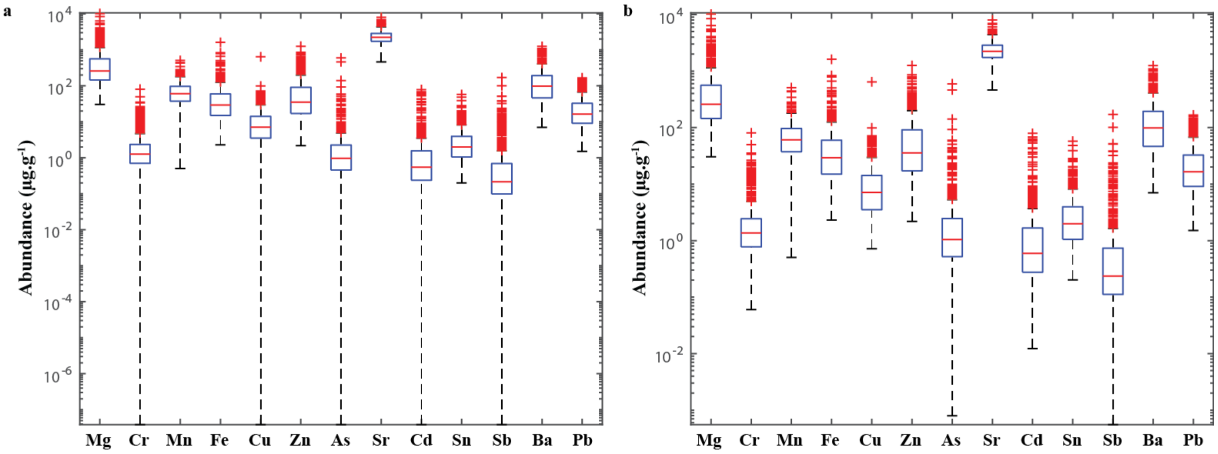


**Figure S1:** Elemental composition of 600 shells of *Shinkailepas tollmanni* encapsulated larvae. **a:** All data, including measurements below the limit of detection. **b:** Same data excluding values below the limit of detection (for Cr, Cu, As, Cd, and Sb).

Parametric statistical comparison between sites was hampered by lack of normality of the data distributions, which is a requirement for MANOVA, as well as homoscedasticity. Data transformation (log and other methods such as cubic root or fourth root) failed to normalize the data, as observed by other studies measuring low-concentration elements (*e.g.*, Gillanders et al., 2001; de Vries et al., 2005). This step was therefore discarded in favor of multivariate classification, described in the main manuscript.

**Supplementary Information S2**

**Laserogram of acquisition of composition of encapsulated larval shell**

A typical laserogram obtained from an Agilent 8900 ICP-MS Triple Quad by the ablation of a *Shinkailepas tollmanni* larval shell with a femtosecond laser ablation system (Novalase SA) is shown on **Figure S2**. We present here all the elements used for the classification models, as well as Ca which was used as the internal standard. For each shell ablated, another ablation was performed immediately nearby to analyse the tape on which the samples were deposited. Only a small signal is visible from the tape for Mg and Fe, in significantly lower abundance than in the shell, therefore strongly limiting the potential impact of contamination.

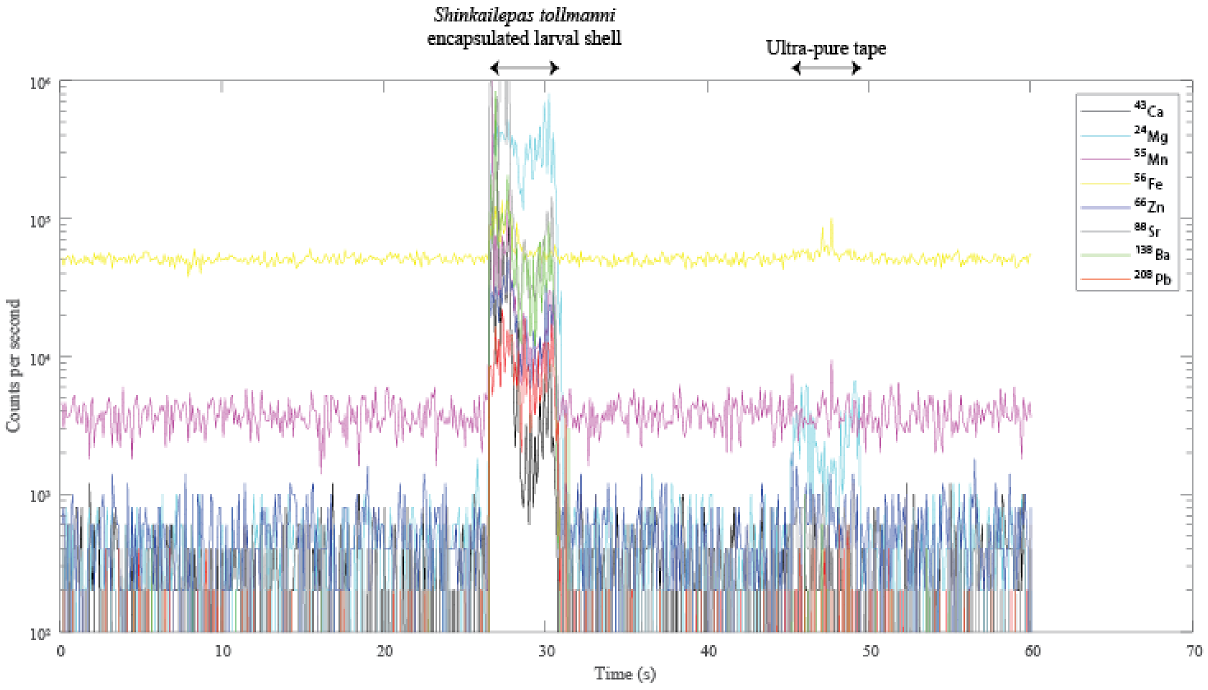


**Figure S2:** Laserogram of a *Shinkailepas tollmanni* encapsulated larval shell and the tape for elements of interest. The positions (in time) of the ablated shell and tape are indicated.

**Supplementary Information S3**

**Best classification models for *Shinkailepas tollmanni* larval shells elemental fingerprinting**

Each model presented in this manuscript refers to the result of a classification method based on several predictors (*i.e.*, variables, here chemical elements). These models can be used as mathematical equations to determine the origin of individual larvae from the composition of its shell.

**Table S3:** Details on models performed for classification.

| **Model** | **Corresponding Figure number** | **Most accurate method** | **Classification success rate** | **Predictors (number)** |
|---|---|---|---|---|
| All sites | Figure 4 | Quadratic Support Vector Machines | 70.0% | Mg, Mn, Fe, Zn, Sr, Ba, Pb (7) |
| Manus/Woodlark | Figure 5a | Trilayered Neural Network | 86.5% | Mn, Ba (2) |
| Sites in East/Woodlark | Figure 5b | Quadratic Support Vector Machines | 77.1% | Mg, Mn, Fe, Zn, Sr, Ba, Pb (7) |
| Areas in East/Woodlark | Figure 5c | Quadratic Support Vector Machines | 75.3% | Mg, Mn, Fe, Zn, Sr, Ba, Pb (7) |
| Sites in Lau | Figure 6a | Linear Discriminant | 97.7% | Mn, Ba (2) |
| Sites in Mangatolo | Figure 6b | Medium Gaussian Support Vector Machines | 98.6% | Fe, Ba (2) |
| Sites in Futuna | Figure 6c | Medium Gaussian Support Vector Machines | 100% | Mn, Ba (2) |

*Short explanations on used methods*

To classify data, a support vector machine (SVM) finds the best hyperplane separating all data points of one class (here, site/area) from those of other classes.

Neural networks learn how to classify data using interconnected nodes in a layered structure that resembles a human brain with neurons. Each node transforms input data using single operators and sends the result to the next layer. Each layer learns from the previous one (the first one being the initial training data) according to how its individual nodes are connected and by the weights of those connections.

Discriminant analysis assumes that different classes present data based on different multivariate Gaussian distributions, and estimates the parameters of that distribution for each class. Linear discriminant analysis is also known as the Fisher discriminant.

The best model to classify Lau Basin sites uses Linear Discriminant Analysis, although the distribution of data on both kept variables (Mn and Ba) do not exhibit a normal distribution (one-sample Kolmogorov-Smirnov tests, 5% significance level), also visible on the scatter plot of Figure 6 of the manuscript. However, the distribution of the data from this area presents some overlap between the sites, which tend to reduce the quality of the discrimination of the SVM method. For comparison, Medium Gaussian SVM reached a 97.0% classification success rate on this dataset. Although the difference is minimal, we elected to keep the Linear Discriminant-based model that present better results.

**Supplementary Information S4**

**Matlab scripts and functions for randomization.**

This document is composed of ten distinct files, to save each as .m files.

The first one is the script that computes the randomization process of the data to be checked using the user-defined trained classifier, modified from White & Ruttenberg (2007) which was restricted to linear discriminant analysis. This modified version allows running this procedure on any type of classification model saved from the *ClassificationLearner* app from Matlab. To do so, select a trainer classifier from the app, click on “Generate function” (top right corner of the app on Matlab version R2022a), and save it on your computer. The name of that function is to be indicated in the script where indicated.

The code by White & Ruttenberg (2007) used site numbers instead of site names. Since we used site names to train the model, we had to convert site names to site numbers, and site numbers to site names to call the trained classifier. Two functions have been written to perform this task and should be modified according to your data. These two functions are given after the script described above.

The following functions correspond to the trained classifiers generated from our dataset for each of the seven models presented in our manuscript.

***Script for randomization***

```
%This script generates a null expectation for the jack-knife
%reclassification success of a discriminant function analysis by
%randomizing user-provided data

%Copyright 2005 J. Wilson White
%Dept. of Ecology, Evolution, and Marine Biology
%University of California, Santa Barbara, CA 93106
%

%version: last modified March 8, 2007



%Modified by Vincent Mouchi on September 13, 2022


% The initial code by J. Wilson White required numbers as sites. Since we
% used site names to train the model, we had to convert site names to site
% numbers, and site numbers to site names to call the trained classifier.
% Two functions have been written to perform this task (available on
% separate files).

%Output: Vector 'null' containing (in order):
%        Mean of randomized distribution (null(1))
%        Standard deviation of randomized distribution (null(2))
%        Median of randomized distribution (null(3))
%        95% quantile of randomized distribution (null(4))
%        5% quantile of randomized distribution  (null(5))
%        Actual jackknife value (null(6))
%        One-tailed P-value (alpha = 0.05) for actual jackknife value
%        (p(7))



%% To edit
Data=Data_all_sites; % User matrix of data
Group_names=Site_names_all;  % Class names (corresponding to those used for the
trained model)
runs=1000;   % Number of bootstraps desired

% Change also the name of the trained classifier at lines 78 & 108
% Change also the name of the function to convert numbers in sites or
% regions at lines 48, 86 & 112 for class to number, and 78 & 108 for number to
class

%% End edit


nbind=size(Data,1);
Groupid = tollmanni_site2number(Group_names);    %vector Groupid holds group
labels

%%
%Data = Data;                %matrix Data holds raw data
%vars = length(D(1,2:end));      %vars = #columns in Data
g = length(Groupid);            %g = #rows in Data
```

```
rdist=[];

%Begin randomization
disp('Step 1: randomization')
for z=1:runs
    disp(sprintf('Number of iteration: %d',z))
    %Assemble randomized dataset:
    newgroups = randperm(g);
    for b = 1:g %for each sample
        rgroups(b) = Groupid(newgroups(b)); %select random group ID
    end    %end for each sample


%the following loop performs the jackknife
for i=1:nbind %begin jackknifing
    data2=Data;
    group2=rgroups;
    group2(i)=[]; %delete the group ID to be jackknifed out
    data2(i,:)=[]; %delete the data to be jackknifed out
    sample=Data(i,:);

    % Modification here by Vincent Mouchi for classification method %
    [trainedClassifier, validationAccuracy] =
trainClassifierAllSites_70_QuadraticSVM(data2, tollmanni_number2site(group2));
    class(i) = trainedClassifier.predictFcn(sample);
    %class(i)=classify(sample,data2,group2,'linear'); %Matlab's DFA function
    %disp(i)

end %end jackknifing


class_id=tollmanni_site2number(class);

q = 0;  %q is counter for DFA classification success
q = sum(class_id == rgroups) %tabulate correct classifications

dis=q/length(Data(:,1))    %calculate jackknife success rate

rdist(z)=dis;%store success rate in vector 'rdist'
%clear class
end %end randomizations


%% Now calculate observed jackknife success rate for the data:
disp('Step 2: data')
for i=1:nbind
    data2=Data;
    group2=Groupid;
    group2(i)=[];
    data2(i, :) = [];
    sample=Data(i,:);

    % Modification here by Vincent Mouchi for classification method %
    [trainedClassifier, validationAccuracy] =
trainClassifierAllSites_70_QuadraticSVM(data2, tollmanni_number2site(group2));
    class(i) = trainedClassifier.predictFcn(sample);
end
```

```
class_num=tollmanni_site2number(class);

q = 0;
q = sum(class_num == Groupid) %tabulate correct classifications
Uservalue=q/length(Groupid)    %calculate observed jackknife success rate

%% Calculate Pvalue

rdist = sort(rdist); %sort rdist in ascending order

i = 1;
while rdist(i) < Uservalue && i < runs %determine where actual jackknife value
lies in null distribution
    i = i + 1;
end

Pvalue = 1-(i)/runs; %one-tailed P-value based on position of actual jackknife
value in null distribution
if Pvalue == 0
    Pvalue = 1/runs;  %the smallest possible P-value provided is 1/runs
end



null=[mean(rdist) std(rdist) quantile(rdist,0.5) quantile(rdist,0.95)
quantile(rdist,0.05) Uservalue Pvalue];
%Report results
fprintf('\n','Results:','\n')
Mean=null(1)
sd = null(2)
Median = null(3)
Upper95quantile = null(4)
Lower95quantile = null(5)
Observed = Uservalue
P = Pvalue
```

***Function to convert site names in numbers***

```
function site_number=tollmanni_site2number(site_name)
% Convert the site name from tollmanni data to site number
% site_number=tollmanni_site2number(site_name)
% Input: site_name: cell, name of site
% Output: site_number: integer

for i=1:length(site_name)
if strcmp(site_name(i),'ABE')
    site_number(i)=1;
else
    if strcmp(site_name(i),'Fenway')
        site_number(i)=2;
    else
        if strcmp(site_name(i),'Futuna North')
            site_number(i)=3;
        else
            if strcmp(site_name(i),'Futuna South')
                site_number(i)=4;
            else
                if strcmp(site_name(i),'Mangatolo North')
                    site_number(i)=5;
                else
                    if strcmp(site_name(i),'Mangatolo South')
                        site_number(i)=6;
                    else
                        if strcmp(site_name(i),'North_Su')
                            site_number(i)=7;
                        else
                            if strcmp(site_name(i),'Phoenix')
                                site_number(i)=8;
                            else
                                if strcmp(site_name(i),'Roman_Ruins')
                                    site_number(i)=9;
                                else
                                    if strcmp(site_name(i),'Scala')
                                        site_number(i)=10;
                                    else
                                        if strcmp(site_name(i),'Solwara')
                                            site_number(i)=11;
                                        else
                                            if strcmp(site_name(i),'South_Su')
                                                site_number(i)=12;
                                            else
                                                if strcmp(site_name(i),'Tow cam')
                                                    site_number(i)=13;
                                                else
                                                    if strcmp(site_name(i),'Tui Malila')
                                                        site_number(i)=14;
                                                    end
                                                end
                                            end
                                        end
                                    end
                                end
                            end
                        end
                    end
                end
            end
        end
    end
end
end
end
```

***Function to convert site numbers in names***

```
function site_name=tollmanni_number2site(site_number)
% Convert the site number from tollmanni data to site name
% site_name=tollmanni_number2site(site_number)
% Output: site_name: cell, name of site
% Input: site_number: integer

for i=1:length(site_number)
if site_number(i)==1
    site_name{i}='ABE';
else
    if site_number(i)==2
        site_name{i}='Fenway';
    else
        if site_number(i)==3
            site_name{i}='Futuna North';
        else
            if site_number(i)==4
                site_name{i}='Futuna South';
            else
                if site_number(i)==5
                    site_name{i}='Mangatolo North';
                else
                    if site_number(i)==6
                        site_name{i}='Mangatolo South';
                    else
                        if site_number(i)==7
                            site_name{i}='North_Su';
                        else
                            if site_number(i)==8
                                site_name{i}='Phoenix';
                            else
                                if site_number(i)==9
                                    site_name{i}='Roman_Ruins';
                                else
                                    if site_number(i)==10
                                        site_name{i}='Scala';
                                    else
                                        if site_number(i)==11
                                            site_name{i}='Solwara';
                                        else
                                            if site_number(i)==12
                                                site_name{i}='South_Su';
                                            else
                                                if site_number(i)==13
                                                    site_name{i}='Tow cam';
                                                else
                                                    if site_number(i)==14
                                                        site_name{i}='Tui Malila';
                                                    end
                                                end
                                            end
                                        end
                                    end
                                end
                            end
                        end
                    end
                end
```

```
                    end
                end
            end
        end
    end
    end
    end
```

***Trained classifier for all sites by Quadratic SVM***

```
function [trainedClassifier, validationAccuracy] =
trainClassifierAllSites_70_QuadraticSVM(trainingData, responseData)
% [trainedClassifier, validationAccuracy] = trainClassifier(trainingData,
% responseData)
% Returns a trained classifier and its accuracy. This code recreates the
% classification model trained in Classification Learner app. Use the
% generated code to automate training the same model with new data, or to
% learn how to programmatically train models.
%
%  Input:
%      trainingData: A matrix with the same number of columns and data type
%       as the matrix imported into the app.
%
%      responseData: A vector with the same data type as the vector
%       imported into the app. The length of responseData and the number of
%       rows of trainingData must be equal.
%
%  Output:
%      trainedClassifier: A struct containing the trained classifier. The
%       struct contains various fields with information about the trained
%       classifier.
%
%      trainedClassifier.predictFcn: A function to make predictions on new
%       data.
%
%      validationAccuracy: A double containing the accuracy as a
%       percentage. In the app, the Models pane displays this overall
%       accuracy score for each model.
%
% Use the code to train the model with new data. To retrain your
% classifier, call the function from the command line with your original
% data or new data as the input arguments trainingData and responseData.
%
% For example, to retrain a classifier trained with the original data set T
% and response Y, enter:
%   [trainedClassifier, validationAccuracy] = trainClassifier(T, Y)
%
% To make predictions with the returned 'trainedClassifier' on new data T2,
% use
%   yfit = trainedClassifier.predictFcn(T2)
%
% T2 must be a matrix containing only the predictor columns used for
% training. For details, enter:
%   trainedClassifier.HowToPredict

warning('off','all')  % Do not display warnings (for trainClassifier)


% Auto-generated by MATLAB on 09-Sep-2022 16:42:26


% Extract predictors and response
% This code processes the data into the right shape for training the
% model.
% Convert input to table
```

```
inputTable = array2table(trainingData, 'VariableNames', {'column_1', 'column_2',
'column_3', 'column_4', 'column_5', 'column_6', 'column_7', 'column_8',
'column_9', 'column_10', 'column_11', 'column_12', 'column_13'});

predictorNames = {'column_1', 'column_2', 'column_3', 'column_4', 'column_5',
'column_6', 'column_7', 'column_8', 'column_9', 'column_10', 'column_11',
'column_12', 'column_13'};
predictors = inputTable(:, predictorNames);
response = responseData;
isCategoricalPredictor = [false, false, false, false, false, false, false, false,
false, false, false, false, false];

% Data transformation: Select subset of the features
% This code selects the same subset of features as were used in the app.
includedPredictorNames = predictors.Properties.VariableNames([true false true true
false true false true false false false true true]);
predictors = predictors(:,includedPredictorNames);
isCategoricalPredictor = isCategoricalPredictor([true false true true false true
false true false false false true true]);

% Train a classifier
% This code specifies all the classifier options and trains the classifier.
template = templateSVM(...
    'KernelFunction', 'polynomial', ...
    'PolynomialOrder', 2, ...
    'KernelScale', 'auto', ...
    'BoxConstraint', 1, ...
    'Standardize', false);
classificationSVM = fitcecoc(...
    predictors, ...
    response, ...
    'Learners', template, ...
    'Coding', 'onevsone', ...
    'ClassNames', {'ABE'; 'Fenway'; 'Futuna North'; 'Futuna South'; 'Mangatolo
North'; 'Mangatolo South'; 'North_Su'; 'Phoenix'; 'Roman_Ruins'; 'Scala';
'Solwara'; 'South_Su'; 'Tow cam'; 'Tui Malila'});

% Create the result struct with predict function
predictorExtractionFcn = @(x) array2table(x, 'VariableNames', predictorNames);
featureSelectionFcn = @(x) x(:,includedPredictorNames);
svmPredictFcn = @(x) predict(classificationSVM, x);
trainedClassifier.predictFcn = @(x)
svmPredictFcn(featureSelectionFcn(predictorExtractionFcn(x)));

% Add additional fields to the result struct
trainedClassifier.ClassificationSVM = classificationSVM;
trainedClassifier.About = 'This struct is a trained model exported from
Classification Learner R2022a.';
trainedClassifier.HowToPredict = sprintf('To make predictions on a new predictor
column matrix, X, use: \n  yfit = c.predictFcn(X) \nreplacing ''c'' with the name
of the variable that is this struct, e.g. ''trainedModel''. \n \nX must contain
exactly 13 columns because this model was trained using 13 predictors. \nX must
contain only predictor columns in exactly the same order and format as your
training \ndata. Do not include the response column or any columns you did not
import into the app. \n \nFor more information, see <a
href="matlab:helpview(fullfile(docroot, ''stats'', ''stats.map''),
''appclassification_exportmodeltoworkspace'')">How to predict using an exported
model</a>.');
```

```
% Extract predictors and response
% This code processes the data into the right shape for training the
% model.
% Convert input to table
inputTable = array2table(trainingData, 'VariableNames', {'column_1', 'column_2',
'column_3', 'column_4', 'column_5', 'column_6', 'column_7', 'column_8',
'column_9', 'column_10', 'column_11', 'column_12', 'column_13'});

predictorNames = {'column_1', 'column_2', 'column_3', 'column_4', 'column_5',
'column_6', 'column_7', 'column_8', 'column_9', 'column_10', 'column_11',
'column_12', 'column_13'};
predictors = inputTable(:, predictorNames);
response = responseData;
isCategoricalPredictor = [false, false, false, false, false, false, false, false,
false, false, false, false, false];

% Perform cross-validation
partitionedModel = crossval(trainedClassifier.ClassificationSVM, 'KFold', 5);

% Compute validation predictions
[validationPredictions, validationScores] = kfoldPredict(partitionedModel);

% Compute validation accuracy
validationAccuracy = 1 - kfoldLoss(partitionedModel, 'LossFun', 'ClassifError');
```

***Trained classifier for sites from Manus and Woodlark by Trilayered NN***

```
function [trainedClassifier, validationAccuracy] =
trainClassifierManusWoodlark_TrilayeredNN(trainingData, responseData)
% [trainedClassifier, validationAccuracy] =
trainClassifierManusWoodlark_TrilayeredNN(trainingData,
% responseData)
% Returns a trained classifier and its accuracy. This code recreates the
% classification model trained in Classification Learner app. Use the
% generated code to automate training the same model with new data, or to
% learn how to programmatically train models.
%
%  Input:
%      trainingData: A matrix with the same number of columns and data type
%       as the matrix imported into the app.
%
%      responseData: A vector with the same data type as the vector
%       imported into the app. The length of responseData and the number of
%       rows of trainingData must be equal.
%
%  Output:
%      trainedClassifier: A struct containing the trained classifier. The
%       struct contains various fields with information about the trained
%       classifier.
%
%      trainedClassifier.predictFcn: A function to make predictions on new
%       data.
%
%      validationAccuracy: A double containing the accuracy as a
%       percentage. In the app, the Models pane displays this overall
%       accuracy score for each model.
%
% Use the code to train the model with new data. To retrain your
% classifier, call the function from the command line with your original
% data or new data as the input arguments trainingData and responseData.
%
% For example, to retrain a classifier trained with the original data set T
% and response Y, enter:
%   [trainedClassifier, validationAccuracy] = trainClassifier(T, Y)
%
% To make predictions with the returned 'trainedClassifier' on new data T2,
% use
%   yfit = trainedClassifier.predictFcn(T2)
%
% T2 must be a matrix containing only the predictor columns used for
% training. For details, enter:
%   trainedClassifier.HowToPredict

% Auto-generated by MATLAB on 14-Nov-2022 10:45:58

warning('off','all')  % Do not display warnings

% Extract predictors and response
% This code processes the data into the right shape for training the
% model.
% Convert input to table
```

```
inputTable = array2table(trainingData, 'VariableNames', {'column_1', 'column_2',
'column_3', 'column_4', 'column_5', 'column_6', 'column_7', 'column_8',
'column_9', 'column_10', 'column_11', 'column_12', 'column_13'});

predictorNames = {'column_1', 'column_2', 'column_3', 'column_4', 'column_5',
'column_6', 'column_7', 'column_8', 'column_9', 'column_10', 'column_11',
'column_12', 'column_13'};
predictors = inputTable(:, predictorNames);
response = responseData;
isCategoricalPredictor = [false, false, false, false, false, false, false, false,
false, false, false, false, false];

% Data transformation: Select subset of the features
% This code selects the same subset of features as were used in the app.
includedPredictorNames = predictors.Properties.VariableNames([false false true
false false false false false false false false true false]);
predictors = predictors(:,includedPredictorNames);
isCategoricalPredictor = isCategoricalPredictor([false false true false false
false false false false false false true false]);

% Train a classifier
% This code specifies all the classifier options and trains the classifier.
classificationNeuralNetwork = fitcnet(...
    predictors, ...
    response, ...
    'LayerSizes', [10 10 10], ...
    'Activations', 'relu', ...
    'Lambda', 0, ...
    'IterationLimit', 1000, ...
    'Standardize', true, ...
    'ClassNames', {'Fenway'; 'North_Su'; 'Roman_Ruins'; 'Scala'; 'Solwara';
'South_Su'});

% Create the result struct with predict function
predictorExtractionFcn = @(x) array2table(x, 'VariableNames', predictorNames);
featureSelectionFcn = @(x) x(:,includedPredictorNames);
neuralNetworkPredictFcn = @(x) predict(classificationNeuralNetwork, x);
trainedClassifier.predictFcn = @(x)
neuralNetworkPredictFcn(featureSelectionFcn(predictorExtractionFcn(x)));

% Add additional fields to the result struct
trainedClassifier.ClassificationNeuralNetwork = classificationNeuralNetwork;
trainedClassifier.About = 'This struct is a trained model exported from
Classification Learner R2022a.';
trainedClassifier.HowToPredict = sprintf('To make predictions on a new predictor
column matrix, X, use: \n  yfit = c.predictFcn(X) \nreplacing ''c'' with the name
of the variable that is this struct, e.g. ''trainedModel''. \n \nX must contain
exactly 13 columns because this model was trained using 13 predictors. \nX must
contain only predictor columns in exactly the same order and format as your
training \ndata. Do not include the response column or any columns you did not
import into the app. \n \nFor more information, see <a
href="matlab:helpview(fullfile(docroot, ''stats'', ''stats.map''),
''appclassification_exportmodeltoworkspace'')">How to predict using an exported
model</a>.');

% Extract predictors and response
% This code processes the data into the right shape for training the
% model.
% Convert input to table
```

```
inputTable = array2table(trainingData, 'VariableNames', {'column_1', 'column_2',
'column_3', 'column_4', 'column_5', 'column_6', 'column_7', 'column_8',
'column_9', 'column_10', 'column_11', 'column_12', 'column_13'});

predictorNames = {'column_1', 'column_2', 'column_3', 'column_4', 'column_5',
'column_6', 'column_7', 'column_8', 'column_9', 'column_10', 'column_11',
'column_12', 'column_13'};
predictors = inputTable(:, predictorNames);
response = responseData;
isCategoricalPredictor = [false, false, false, false, false, false, false, false,
false, false, false, false, false];

% Perform cross-validation
partitionedModel = crossval(trainedClassifier.ClassificationNeuralNetwork,
'KFold', 5);

% Compute validation predictions
[validationPredictions, validationScores] = kfoldPredict(partitionedModel);

% Compute validation accuracy
validationAccuracy = 1 - kfoldLoss(partitionedModel, 'LossFun', 'ClassifError');
```

***Trained classifier for sites from eastern region and Woodlark by Quadratic SVM***

```
function [trainedClassifier, validationAccuracy] =
trainClassifierSitesEasternWoodlark_77_1_QuadraticSVM(trainingData, responseData)
% [trainedClassifier, validationAccuracy] =
trainClassifierSitesEasternWoodlark_77_1_QuadraticSVM(trainingData,
% responseData)
% Returns a trained classifier and its accuracy. This code recreates the
% classification model trained in Classification Learner app. Use the
% generated code to automate training the same model with new data, or to
% learn how to programmatically train models.
%
%  Input:
%      trainingData: A matrix with the same number of columns and data type
%       as the matrix imported into the app.
%
%      responseData: A vector with the same data type as the vector
%       imported into the app. The length of responseData and the number of
%       rows of trainingData must be equal.
%
%  Output:
%      trainedClassifier: A struct containing the trained classifier. The
%       struct contains various fields with information about the trained
%       classifier.
%
%      trainedClassifier.predictFcn: A function to make predictions on new
%       data.
%
%      validationAccuracy: A double containing the accuracy as a
%       percentage. In the app, the Models pane displays this overall
%       accuracy score for each model.
%
% Use the code to train the model with new data. To retrain your
% classifier, call the function from the command line with your original
% data or new data as the input arguments trainingData and responseData.
%
% For example, to retrain a classifier trained with the original data set T
% and response Y, enter:
%   [trainedClassifier, validationAccuracy] = trainClassifier(T, Y)
%
% To make predictions with the returned 'trainedClassifier' on new data T2,
% use
%   yfit = trainedClassifier.predictFcn(T2)
%
% T2 must be a matrix containing only the predictor columns used for
% training. For details, enter:
%   trainedClassifier.HowToPredict

% Auto-generated by MATLAB on 06-Oct-2022 13:16:32

warning('off','all')  % Do not display warnings

% Extract predictors and response
% This code processes the data into the right shape for training the
% model.
% Convert input to table
```

```
inputTable = array2table(trainingData, 'VariableNames', {'column_1', 'column_2',
'column_3', 'column_4', 'column_5', 'column_6', 'column_7', 'column_8',
'column_9', 'column_10', 'column_11', 'column_12', 'column_13'});

predictorNames = {'column_1', 'column_2', 'column_3', 'column_4', 'column_5',
'column_6', 'column_7', 'column_8', 'column_9', 'column_10', 'column_11',
'column_12', 'column_13'};
predictors = inputTable(:, predictorNames);
response = responseData;
isCategoricalPredictor = [false, false, false, false, false, false, false, false,
false, false, false, false, false];

% Data transformation: Select subset of the features
% This code selects the same subset of features as were used in the app.
includedPredictorNames = predictors.Properties.VariableNames([true false true true
false true false true false false false true true]);
predictors = predictors(:,includedPredictorNames);
isCategoricalPredictor = isCategoricalPredictor([true false true true false true
false true false false false true true]);

% Train a classifier
% This code specifies all the classifier options and trains the classifier.
template = templateSVM(...
    'KernelFunction', 'polynomial', ...
    'PolynomialOrder', 2, ...
    'KernelScale', 'auto', ...
    'BoxConstraint', 1, ...
    'Standardize', true);
classificationSVM = fitcecoc(...
    predictors, ...
    response, ...
    'Learners', template, ...
    'Coding', 'onevsone', ...
    'ClassNames', {'ABE'; 'Futuna North'; 'Futuna South'; 'Mangatolo North';
'Mangatolo South'; 'Phoenix'; 'Scala'; 'Tow cam'; 'Tui Malila'});

% Create the result struct with predict function
predictorExtractionFcn = @(x) array2table(x, 'VariableNames', predictorNames);
featureSelectionFcn = @(x) x(:,includedPredictorNames);
svmPredictFcn = @(x) predict(classificationSVM, x);
trainedClassifier.predictFcn = @(x)
svmPredictFcn(featureSelectionFcn(predictorExtractionFcn(x)));

% Add additional fields to the result struct
trainedClassifier.ClassificationSVM = classificationSVM;
trainedClassifier.About = 'This struct is a trained model exported from
Classification Learner R2022a.';
trainedClassifier.HowToPredict = sprintf('To make predictions on a new predictor
column matrix, X, use: \n  yfit = c.predictFcn(X) \nreplacing ''c'' with the name
of the variable that is this struct, e.g. ''trainedModel''. \n \nX must contain
exactly 13 columns because this model was trained using 13 predictors. \nX must
contain only predictor columns in exactly the same order and format as your
training \ndata. Do not include the response column or any columns you did not
import into the app. \n \nFor more information, see <a
href="matlab:helpview(fullfile(docroot, ''stats'', ''stats.map''),
''appclassification_exportmodeltoworkspace'')">How to predict using an exported
model</a>.');

% Extract predictors and response
```

```
% This code processes the data into the right shape for training the
% model.
% Convert input to table
inputTable = array2table(trainingData, 'VariableNames', {'column_1', 'column_2',
'column_3', 'column_4', 'column_5', 'column_6', 'column_7', 'column_8',
'column_9', 'column_10', 'column_11', 'column_12', 'column_13'});

predictorNames = {'column_1', 'column_2', 'column_3', 'column_4', 'column_5',
'column_6', 'column_7', 'column_8', 'column_9', 'column_10', 'column_11',
'column_12', 'column_13'};
predictors = inputTable(:, predictorNames);
response = responseData;
isCategoricalPredictor = [false, false, false, false, false, false, false, false,
false, false, false, false, false];

% Perform cross-validation
partitionedModel = crossval(trainedClassifier.ClassificationSVM, 'KFold', 5);

% Compute validation predictions
[validationPredictions, validationScores] = kfoldPredict(partitionedModel);

% Compute validation accuracy
validationAccuracy = 1 - kfoldLoss(partitionedModel, 'LossFun', 'ClassifError');
```

***Trained classifier for areas from eastern region and Woodlark by Quadratic SVM***

```
function [trainedClassifier, validationAccuracy] =
trainClassifierAreasEastWoodlark75_3_QuadraticSVM(trainingData, responseData)
% [trainedClassifier, validationAccuracy] =
trainClassifierAreasEastWoodlark75_3_QuadraticSVM(trainingData,
% responseData)
% Returns a trained classifier and its accuracy. This code recreates the
% classification model trained in Classification Learner app. Use the
% generated code to automate training the same model with new data, or to
% learn how to programmatically train models.
%
%  Input:
%      trainingData: A matrix with the same number of columns and data type
%       as the matrix imported into the app.
%
%      responseData: A vector with the same data type as the vector
%       imported into the app. The length of responseData and the number of
%       rows of trainingData must be equal.
%
%  Output:
%      trainedClassifier: A struct containing the trained classifier. The
%       struct contains various fields with information about the trained
%       classifier.
%
%      trainedClassifier.predictFcn: A function to make predictions on new
%       data.
%
%      validationAccuracy: A double containing the accuracy as a
%       percentage. In the app, the Models pane displays this overall
%       accuracy score for each model.
%
% Use the code to train the model with new data. To retrain your
% classifier, call the function from the command line with your original
% data or new data as the input arguments trainingData and responseData.
%
% For example, to retrain a classifier trained with the original data set T
% and response Y, enter:
%   [trainedClassifier, validationAccuracy] = trainClassifier(T, Y)
%
% To make predictions with the returned 'trainedClassifier' on new data T2,
% use
%   yfit = trainedClassifier.predictFcn(T2)
%
% T2 must be a matrix containing only the predictor columns used for
% training. For details, enter:
%   trainedClassifier.HowToPredict

% Auto-generated by MATLAB on 06-Oct-2022 11:34:01
warning('off','all')  % Do not display warnings


% Extract predictors and response
% This code processes the data into the right shape for training the
% model.
% Convert input to table
```

```
inputTable = array2table(trainingData, 'VariableNames', {'column_1', 'column_2',
'column_3', 'column_4', 'column_5', 'column_6', 'column_7', 'column_8',
'column_9', 'column_10', 'column_11', 'column_12', 'column_13'});

predictorNames = {'column_1', 'column_2', 'column_3', 'column_4', 'column_5',
'column_6', 'column_7', 'column_8', 'column_9', 'column_10', 'column_11',
'column_12', 'column_13'};
predictors = inputTable(:, predictorNames);
response = responseData;
isCategoricalPredictor = [false, false, false, false, false, false, false, false,
false, false, false, false, false];

% Data transformation: Select subset of the features
% This code selects the same subset of features as were used in the app.
includedPredictorNames = predictors.Properties.VariableNames([true false true
false false true false true false false false true true]);
predictors = predictors(:,includedPredictorNames);
isCategoricalPredictor = isCategoricalPredictor([true false true false false true
false true false false false true true]);

% Train a classifier
% This code specifies all the classifier options and trains the classifier.
template = templateSVM(...
    'KernelFunction', 'polynomial', ...
    'PolynomialOrder', 2, ...
    'KernelScale', 'auto', ...
    'BoxConstraint', 1, ...
    'Standardize', true);
classificationSVM = fitcecoc(...
    predictors, ...
    response, ...
    'Learners', template, ...
    'Coding', 'onevsone', ...
    'ClassNames', {'Futuna'; 'Lau'; 'Mangatolo'; 'North Fiji'; 'Woodlark'});

% Create the result struct with predict function
predictorExtractionFcn = @(x) array2table(x, 'VariableNames', predictorNames);
featureSelectionFcn = @(x) x(:,includedPredictorNames);
svmPredictFcn = @(x) predict(classificationSVM, x);
trainedClassifier.predictFcn = @(x)
svmPredictFcn(featureSelectionFcn(predictorExtractionFcn(x)));

% Add additional fields to the result struct
trainedClassifier.ClassificationSVM = classificationSVM;
trainedClassifier.About = 'This struct is a trained model exported from
Classification Learner R2022a.';
trainedClassifier.HowToPredict = sprintf('To make predictions on a new predictor
column matrix, X, use: \n  yfit = c.predictFcn(X) \nreplacing ''c'' with the name
of the variable that is this struct, e.g. ''trainedModel''. \n \nX must contain
exactly 13 columns because this model was trained using 13 predictors. \nX must
contain only predictor columns in exactly the same order and format as your
training \ndata. Do not include the response column or any columns you did not
import into the app. \n \nFor more information, see <a
href="matlab:helpview(fullfile(docroot, ''stats'', ''stats.map''),
''appclassification_exportmodeltoworkspace'')">How to predict using an exported
model</a>.');

% Extract predictors and response
% This code processes the data into the right shape for training the
```

```
% model.
% Convert input to table
inputTable = array2table(trainingData, 'VariableNames', {'column_1', 'column_2',
'column_3', 'column_4', 'column_5', 'column_6', 'column_7', 'column_8',
'column_9', 'column_10', 'column_11', 'column_12', 'column_13'});

predictorNames = {'column_1', 'column_2', 'column_3', 'column_4', 'column_5',
'column_6', 'column_7', 'column_8', 'column_9', 'column_10', 'column_11',
'column_12', 'column_13'};
predictors = inputTable(:, predictorNames);
response = responseData;
isCategoricalPredictor = [false, false, false, false, false, false, false, false,
false, false, false, false, false];

% Perform cross-validation
partitionedModel = crossval(trainedClassifier.ClassificationSVM, 'KFold', 5);

% Compute validation predictions
[validationPredictions, validationScores] = kfoldPredict(partitionedModel);

% Compute validation accuracy
validationAccuracy = 1 - kfoldLoss(partitionedModel, 'LossFun', 'ClassifError');
```

***Trained classifier for sites from Lau by LDA***

```
function [trainedClassifier, validationAccuracy] =
trainClassifierLau_97_7_LinearDiscriminant2(trainingData, responseData)
% [trainedClassifier, validationAccuracy] =
trainClassifierLau_97_7_LinearDiscriminant2(trainingData,
% responseData)
% Returns a trained classifier and its accuracy. This code recreates the
% classification model trained in Classification Learner app. Use the
% generated code to automate training the same model with new data, or to
% learn how to programmatically train models.
%
%  Input:
%      trainingData: A matrix with the same number of columns and data type
%       as the matrix imported into the app.
%
%      responseData: A vector with the same data type as the vector
%       imported into the app. The length of responseData and the number of
%       rows of trainingData must be equal.
%
%  Output:
%      trainedClassifier: A struct containing the trained classifier. The
%       struct contains various fields with information about the trained
%       classifier.
%
%      trainedClassifier.predictFcn: A function to make predictions on new
%       data.
%
%      validationAccuracy: A double containing the accuracy as a
%       percentage. In the app, the Models pane displays this overall
%       accuracy score for each model.
%
% Use the code to train the model with new data. To retrain your
% classifier, call the function from the command line with your original
% data or new data as the input arguments trainingData and responseData.
%
% For example, to retrain a classifier trained with the original data set T
% and response Y, enter:
%   [trainedClassifier, validationAccuracy] = trainClassifier(T, Y)
%
% To make predictions with the returned 'trainedClassifier' on new data T2,
% use
%   yfit = trainedClassifier.predictFcn(T2)
%
% T2 must be a matrix containing only the predictor columns used for
% training. For details, enter:
%   trainedClassifier.HowToPredict

% Auto-generated by MATLAB on 15-Nov-2022 11:12:35

warning('off','all')  % Do not display warnings

% Extract predictors and response
% This code processes the data into the right shape for training the
% model.
% Convert input to table
```

```
inputTable = array2table(trainingData, 'VariableNames', {'column_1', 'column_2',
'column_3', 'column_4', 'column_5', 'column_6', 'column_7', 'column_8',
'column_9', 'column_10', 'column_11', 'column_12', 'column_13'});

predictorNames = {'column_1', 'column_2', 'column_3', 'column_4', 'column_5',
'column_6', 'column_7', 'column_8', 'column_9', 'column_10', 'column_11',
'column_12', 'column_13'};
predictors = inputTable(:, predictorNames);
response = responseData;
isCategoricalPredictor = [false, false, false, false, false, false, false, false,
false, false, false, false, false];

% Data transformation: Select subset of the features
% This code selects the same subset of features as were used in the app.
includedPredictorNames = predictors.Properties.VariableNames([false false true
false false false false false false false false true false]);
predictors = predictors(:,includedPredictorNames);
isCategoricalPredictor = isCategoricalPredictor([false false true false false
false false false false false false true false]);

% Train a classifier
% This code specifies all the classifier options and trains the classifier.
classificationDiscriminant = fitcdiscr(...
    predictors, ...
    response, ...
    'DiscrimType', 'linear', ...
    'Gamma', 0, ...
    'FillCoeffs', 'off', ...
    'ClassNames', {'ABE'; 'Tow cam'; 'Tui Malila'});

% Create the result struct with predict function
predictorExtractionFcn = @(x) array2table(x, 'VariableNames', predictorNames);
featureSelectionFcn = @(x) x(:,includedPredictorNames);
discriminantPredictFcn = @(x) predict(classificationDiscriminant, x);
trainedClassifier.predictFcn = @(x)
discriminantPredictFcn(featureSelectionFcn(predictorExtractionFcn(x)));

% Add additional fields to the result struct
trainedClassifier.ClassificationDiscriminant = classificationDiscriminant;
trainedClassifier.About = 'This struct is a trained model exported from
Classification Learner R2022a.';
trainedClassifier.HowToPredict = sprintf('To make predictions on a new predictor
column matrix, X, use: \n  yfit = c.predictFcn(X) \nreplacing ''c'' with the name
of the variable that is this struct, e.g. ''trainedModel''. \n \nX must contain
exactly 13 columns because this model was trained using 13 predictors. \nX must
contain only predictor columns in exactly the same order and format as your
training \ndata. Do not include the response column or any columns you did not
import into the app. \n \nFor more information, see <a
href="matlab:helpview(fullfile(docroot, ''stats'', ''stats.map''),
''appclassification_exportmodeltoworkspace'')">How to predict using an exported
model</a>.');

% Extract predictors and response
% This code processes the data into the right shape for training the
% model.
% Convert input to table
inputTable = array2table(trainingData, 'VariableNames', {'column_1', 'column_2',
'column_3', 'column_4', 'column_5', 'column_6', 'column_7', 'column_8',
'column_9', 'column_10', 'column_11', 'column_12', 'column_13'});
```

```
predictorNames = {'column_1', 'column_2', 'column_3', 'column_4', 'column_5',
'column_6', 'column_7', 'column_8', 'column_9', 'column_10', 'column_11',
'column_12', 'column_13'};
predictors = inputTable(:, predictorNames);
response = responseData;
isCategoricalPredictor = [false, false, false, false, false, false, false, false,
false, false, false, false, false];

% Perform cross-validation
partitionedModel = crossval(trainedClassifier.ClassificationDiscriminant, 'KFold',
5);

% Compute validation predictions
[validationPredictions, validationScores] = kfoldPredict(partitionedModel);

% Compute validation accuracy
validationAccuracy = 1 - kfoldLoss(partitionedModel, 'LossFun', 'ClassifError');
```

***Trained classifier for sites from Futuna by Medium Gaussian SVM***

```
function [trainedClassifier, validationAccuracy] = 
trainClassifierFutuna_100_MediumGaussianSVM2(trainingData, responseData)
% [trainedClassifier, validationAccuracy] = 
trainClassifierFutuna_100_MediumGaussianSVM2(trainingData,
% responseData)
% Returns a trained classifier and its accuracy. This code recreates the
% classification model trained in Classification Learner app. Use the
% generated code to automate training the same model with new data, or to
% learn how to programmatically train models.
%
%  Input:
%      trainingData: A matrix with the same number of columns and data type
%       as the matrix imported into the app.
%
%      responseData: A vector with the same data type as the vector
%       imported into the app. The length of responseData and the number of
%       rows of trainingData must be equal.
%
%  Output:
%      trainedClassifier: A struct containing the trained classifier. The
%       struct contains various fields with information about the trained
%       classifier.
%
%      trainedClassifier.predictFcn: A function to make predictions on new
%       data.
%
%      validationAccuracy: A double containing the accuracy as a
%       percentage. In the app, the Models pane displays this overall
%       accuracy score for each model.
%
% Use the code to train the model with new data. To retrain your
% classifier, call the function from the command line with your original
% data or new data as the input arguments trainingData and responseData.
%
% For example, to retrain a classifier trained with the original data set T
% and response Y, enter:
%   [trainedClassifier, validationAccuracy] = trainClassifier(T, Y)
%
% To make predictions with the returned 'trainedClassifier' on new data T2,
% use
%   yfit = trainedClassifier.predictFcn(T2)
%
% T2 must be a matrix containing only the predictor columns used for
% training. For details, enter:
%   trainedClassifier.HowToPredict

% Auto-generated by MATLAB on 15-Nov-2022 12:21:34

warning('off','all')  % Do not display warnings

% Extract predictors and response
% This code processes the data into the right shape for training the
% model.
% Convert input to table
```

```
inputTable = array2table(trainingData, 'VariableNames', {'column_1', 'column_2',
'column_3', 'column_4', 'column_5', 'column_6', 'column_7', 'column_8',
'column_9', 'column_10', 'column_11', 'column_12', 'column_13'});

predictorNames = {'column_1', 'column_2', 'column_3', 'column_4', 'column_5',
'column_6', 'column_7', 'column_8', 'column_9', 'column_10', 'column_11',
'column_12', 'column_13'};
predictors = inputTable(:, predictorNames);
response = responseData;
isCategoricalPredictor = [false, false, false, false, false, false, false, false,
false, false, false, false, false];

% Data transformation: Select subset of the features
% This code selects the same subset of features as were used in the app.
includedPredictorNames = predictors.Properties.VariableNames([false false true
false false false false false false false false true false]);
predictors = predictors(:,includedPredictorNames);
isCategoricalPredictor = isCategoricalPredictor([false false true false false
false false false false false false true false]);

% Train a classifier
% This code specifies all the classifier options and trains the classifier.
classificationSVM = fitcsvm(...
    predictors, ...
    response, ...
    'KernelFunction', 'gaussian', ...
    'PolynomialOrder', [], ...
    'KernelScale', 1.4, ...
    'BoxConstraint', 1, ...
    'Standardize', true, ...
    'ClassNames', {'Futuna North'; 'Futuna South'});

% Create the result struct with predict function
predictorExtractionFcn = @(x) array2table(x, 'VariableNames', predictorNames);
featureSelectionFcn = @(x) x(:,includedPredictorNames);
svmPredictFcn = @(x) predict(classificationSVM, x);
trainedClassifier.predictFcn = @(x)
svmPredictFcn(featureSelectionFcn(predictorExtractionFcn(x)));

% Add additional fields to the result struct
trainedClassifier.ClassificationSVM = classificationSVM;
trainedClassifier.About = 'This struct is a trained model exported from
Classification Learner R2022a.';
trainedClassifier.HowToPredict = sprintf('To make predictions on a new predictor
column matrix, X, use: \n  yfit = c.predictFcn(X) \nreplacing ''c'' with the name
of the variable that is this struct, e.g. ''trainedModel''. \n \nX must contain
exactly 13 columns because this model was trained using 13 predictors. \nX must
contain only predictor columns in exactly the same order and format as your
training \ndata. Do not include the response column or any columns you did not
import into the app. \n \nFor more information, see <a
href="matlab:helpview(fullfile(docroot, ''stats'', ''stats.map''),
''appclassification_exportmodeltoworkspace'')">How to predict using an exported
model</a>.');

% Extract predictors and response
% This code processes the data into the right shape for training the
% model.
% Convert input to table
```

```
inputTable = array2table(trainingData, 'VariableNames', {'column_1', 'column_2',
'column_3', 'column_4', 'column_5', 'column_6', 'column_7', 'column_8',
'column_9', 'column_10', 'column_11', 'column_12', 'column_13'});

predictorNames = {'column_1', 'column_2', 'column_3', 'column_4', 'column_5',
'column_6', 'column_7', 'column_8', 'column_9', 'column_10', 'column_11',
'column_12', 'column_13'};
predictors = inputTable(:, predictorNames);
response = responseData;
isCategoricalPredictor = [false, false, false, false, false, false, false, false,
false, false, false, false, false];

% Perform cross-validation
partitionedModel = crossval(trainedClassifier.ClassificationSVM, 'KFold', 5);

% Compute validation predictions
[validationPredictions, validationScores] = kfoldPredict(partitionedModel);

% Compute validation accuracy
validationAccuracy = 1 - kfoldLoss(partitionedModel, 'LossFun', 'ClassifError');
```

***Trained classifier for sites from Mangatolo by SVM***

```
function [trainedClassifier, validationAccuracy] =
trainClassifier_Mangatolo_SVM_98_6(trainingData, responseData)
% [trainedClassifier, validationAccuracy] = trainClassifier(trainingData,
% responseData)
% Returns a trained classifier and its accuracy. This code recreates the
% classification model trained in Classification Learner app. Use the
% generated code to automate training the same model with new data, or to
% learn how to programmatically train models.
%
%  Input:
%      trainingData: A matrix with the same number of columns and data type
%       as the matrix imported into the app.
%
%      responseData: A vector with the same data type as the vector
%       imported into the app. The length of responseData and the number of
%       rows of trainingData must be equal.
%
%
%  Output:
%      trainedClassifier: A struct containing the trained classifier. The
%       struct contains various fields with information about the trained
%       classifier.
%
%      trainedClassifier.predictFcn: A function to make predictions on new
%       data.
%
%      validationAccuracy: A double representing the validation accuracy as
%       a percentage. In the app, the Models pane displays the validation
%       accuracy for each model.
%
% Use the code to train the model with new data. To retrain your
% classifier, call the function from the command line with your original
% data or new data as the input arguments trainingData and responseData.
%
% For example, to retrain a classifier trained with the original data set T
% and response Y, enter:
%   [trainedClassifier, validationAccuracy] = trainClassifier(T, Y)
%
% To make predictions with the returned 'trainedClassifier' on new data T2,
% use
%   [yfit,scores] = trainedClassifier.predictFcn(T2)
%
% T2 must be a matrix containing only the predictor columns used for
% training. For details, enter:
%   trainedClassifier.HowToPredict

% Auto-generated by MATLAB on 01-Apr-2023 12:02:11


% Extract predictors and response
% This code processes the data into the right shape for training the
% model.
% Convert input to table
inputTable = array2table(trainingData, 'VariableNames', {'column_1', 'column_2',
'column_3', 'column_4', 'column_5', 'column_6', 'column_7', 'column_8',
'column_9', 'column_10', 'column_11', 'column_12', 'column_13'});
```

```
predictorNames = {'column_1', 'column_2', 'column_3', 'column_4', 'column_5', 
'column_6', 'column_7', 'column_8', 'column_9', 'column_10', 'column_11', 
'column_12', 'column_13'};
predictors = inputTable(:, predictorNames);
response = responseData;
isCategoricalPredictor = [false, false, false, false, false, false, false, false, 
false, false, false, false, false];
classNames = {'Mangatolo North'; 'Mangatolo South'};

% Data transformation: Select subset of the features
% This code selects the same subset of features as were used in the app.
includedPredictorNames = predictors.Properties.VariableNames([false false false 
true false false false false false false false true false]);
predictors = predictors(:,includedPredictorNames);
isCategoricalPredictor = isCategoricalPredictor([false false false true false 
false false false false false false true false]);

% Train a classifier
% This code specifies all the classifier options and trains the classifier.
classificationSVM = fitcsvm(...
    predictors, ...
    response, ...
    'KernelFunction', 'gaussian', ...
    'PolynomialOrder', [], ...
    'KernelScale', 1.4, ...
    'BoxConstraint', 1, ...
    'Standardize', true, ...
    'ClassNames', classNames);

% Create the result struct with predict function
predictorExtractionFcn = @(x) array2table(x, 'VariableNames', predictorNames);
featureSelectionFcn = @(x) x(:,includedPredictorNames);
svmPredictFcn = @(x) predict(classificationSVM, x);
trainedClassifier.predictFcn = @(x) 
svmPredictFcn(featureSelectionFcn(predictorExtractionFcn(x)));

% Add additional fields to the result struct
trainedClassifier.ClassificationSVM = classificationSVM;
trainedClassifier.About = 'This struct is a trained model exported from 
Classification Learner R2023a.';
trainedClassifier.HowToPredict = sprintf('To make predictions on a new predictor 
column matrix, X, use: \n  [yfit,scores] = c.predictFcn(X) \nreplacing ''c'' with 
the name of the variable that is this struct, e.g. ''trainedModel''. \n \nX must 
contain exactly 13 columns because this model was trained using 13 predictors. \nX 
must contain only predictor columns in exactly the same order and format as your 
training \ndata. Do not include the response column or any columns you did not 
import into the app. \n \nFor more information, see <a 
href="matlab:helpview(fullfile(docroot, ''stats'', ''stats.map''), 
''appclassification_exportmodeltoworkspace'')">How to predict using an exported 
model</a>.');

% Extract predictors and response
% This code processes the data into the right shape for training the
% model.
% Convert input to table
inputTable = array2table(trainingData, 'VariableNames', {'column_1', 'column_2', 
'column_3', 'column_4', 'column_5', 'column_6', 'column_7', 'column_8', 
'column_9', 'column_10', 'column_11', 'column_12', 'column_13'});
```

```
predictorNames = {'column_1', 'column_2', 'column_3', 'column_4', 'column_5',
'column_6', 'column_7', 'column_8', 'column_9', 'column_10', 'column_11',
'column_12', 'column_13'};
predictors = inputTable(:, predictorNames);
response = responseData;
isCategoricalPredictor = [false, false, false, false, false, false, false, false,
false, false, false, false, false];
classNames = {'Mangatolo North'; 'Mangatolo South'};

% Perform cross-validation
partitionedModel = crossval(trainedClassifier.ClassificationSVM, 'KFold', 5);

% Compute validation predictions
[validationPredictions, validationScores] = kfoldPredict(partitionedModel);

% Compute validation accuracy
validationAccuracy = 1 - kfoldLoss(partitionedModel, 'LossFun', 'ClassifError');
```

**Supplementary Information S5**

**Elemental composition of the habitat water surrounding the *Ifremeria nautilei* communities sampled at hydrothermal-vent sites in the Southwest Pacific**

The data presented here are from measurements of 40 seawater samples collected over the *Ifremeria nautilei* communities collected during the Chubacarc expedition (Hourdez & Jollivet 2019). Note that not all of these samples correspond to sites where *Shinkailepas tollmanni* capsules have been collected. All available measurements were used in the manuscript for the PCA. Only 20 of these samples corresponded to locations where *S. tollmanni* capsules were collected, and were used to compare the elemental composition of seawater with that of the larval shells (see Tables 1 and 2 of the manuscript).

**Table S4 (next page):** Concentration data from habitat water.

| Area | Site Name | Ca (mM/kg) | K (mM/kg) | Mg (mM/kg) | S (mM/kg) | Si (µM/kg) | Fe (µM/kg) | Mn (µM/kg) | Cu (µM/kg) | Zn (µM/kg) | Li (µM/kg) | Al (µM/kg) | B (µM/kg) | Rb (µM/kg) | Sr (µM/kg) | V (µM/kg) | P (µM/kg) | Mo (nM/kg) | U (nM/kg) | Y (nM/kg) |
|---|---|---|---|---|---|---|---|---|---|---|---|---|---|---|---|---|---|---|---|---|
| Lau Basin | Tow Cam | 10.37 | 11.19 | 70.75 | 29.61 | 132.94 | 0.11 | 0.15 | 0.06 | 0.43 | 26.39 | 0.44 | 423.45 | 1.40 | 94.33 | 0.03 | 3.19 | 94.70 | 13.13 | 1.17 |
| Lau Basin | Tow Cam | 10.77 | 11.45 | 68.58 | 29.44 | 289.99 | 0.91 | 2.24 | 0.07 | 0.88 | 33.57 | 0.75 | 408.09 | 1.49 | 94.68 | 0.03 | 3.42 | 93.20 | 12.95 | 0.87 |
| Lau Basin | Tu'i Malila | 12.42 | 12.73 | 56.79 | 28.11 | 716.40 | 4.70 | 15.46 | 0.12 | 2.16 | 48.94 | 0.36 | 400.40 | 3.47 | 99.97 | 0.03 | 2.60 | 85.19 | 11.78 | 1.22 |
| Lau Basin | Tu'i Malila | 12.87 | 13.70 | 60.40 | 28.83 | 995.88 | 1.53 | 18.95 | 0.11 | 1.05 | 58.44 | 1.99 | 417.16 | 4.39 | 103.63 | 0.03 | 3.32 | 79.17 | 11.72 | 1.38 |
| Lau Basin | ABE | 10.08 | 11.06 | 52.63 | 26.98 | 299.00 | 0.93 | 3.27 | 0.03 | 0.97 | 29.08 | 1.21 | 423.75 | 1.53 | 86.31 | 0.03 | 4.58 | 84.33 | 11.49 | 0.95 |
| Mangatolo | Mangatolo Triple Junction | 12.04 | 12.83 | 60.44 | 30.43 | 630.38 | 15.87 | 5.81 | 0.46 | 40.28 | 35.48 | 1.36 | 423.97 | 2.74 | 112.69 | 0.04 | 4.90 | 104.48 | 12.14 | 13.23 |
| Mangatolo | Mangatolo Triple Junction | 12.22 | 13.11 | 60.77 | 30.47 | 1105.35 | 14.14 | 28.73 | 0.78 | 20.94 | 35.35 | 1.32 | 450.78 | 2.82 | 103.84 | 0.03 | 4.57 | 138.28 | 12.33 | 7.81 |
| Futuna | Fatu Kapa - Aster'X | 13.24 | 14.40 | 64.82 | 33.47 | 298.79 | 0.86 | 2.06 | 0.07 | 1.47 | 42.72 | 0.90 | 498.16 | 3.43 | 104.49 | 0.04 | 3.84 | 97.65 | 10.37 | 0.90 |
| Futuna | Fatu Kapa -Stéphanie | 13.41 | 14.03 | 64.80 | 33.33 | 332.17 | 1.86 | 4.19 | 0.10 | 1.32 | 40.52 | 1.09 | 473.12 | 3.02 | 105.03 | 0.04 | 4.46 | 95.95 | 11.49 | 0.81 |
| Futuna | Fatu Kapa -Fati Ufu 7 | 11.70 | 11.54 | 52.19 | 27.42 | 440.68 | 3.27 | 7.18 | 0.08 | 1.20 | 41.26 | 0.99 | 416.43 | 3.22 | 93.21 | 0.03 | 2.94 | 96.72 | 12.45 | 0.93 |
| Futuna | Fatu Kapa -Fati Ufu 9 | 14.24 | 12.45 | 51.71 | 26.84 | 517.81 | 10.28 | 20.26 | 0.04 | 0.75 | 56.27 | 0.72 | 414.13 | 5.52 | 97.39 | 0.03 | 2.87 | 88.29 | 11.55 | 1.07 |
| Futuna | Kulo Lasi | 9.61 | 10.03 | 54.13 | 28.16 | 107.87 | 0.01 | 0.03 | 0.03 | 0.60 | 24.40 | 0.77 | 417.48 | 1.33 | 91.33 | 0.03 | 4.10 | 90.43 | 12.63 | 0.79 |
| North Fiji | Phoenix | 11.23 | 12.18 | 60.48 | 30.00 | 284.37 | 1.13 | 2.11 | 0.01 | 1.64 | 28.58 | 0.69 | 400.36 | 1.56 | 97.67 | 0.03 | 3.33 | 97.30 | 12.42 | 0.92 |
| North Fiji | Phoenix | 11.26 | 11.89 | 60.97 | 30.06 | 212.71 | 0.52 | 1.53 | 0.01 | 0.52 | 26.87 | 0.81 | 421.56 | 1.51 | 97.97 | 0.03 | 3.69 | 98.05 | 13.24 | 0.97 |
| Manus Basin | FenWay | 10.27 | 11.17 | 56.27 | 28.75 | 189.09 | 2.45 | 4.52 | 0.05 | 0.41 | 25.69 | 1.11 | 453.28 | 1.46 | 92.21 | 0.04 | 3.49 | 88.91 | 12.12 | 1.40 |
| Manus Basin | FenWay | 10.03 | 10.89 | 54.55 | 27.17 | 167.42 | 1.99 | 3.63 | 0.11 | 0.45 | 27.13 | 0.77 | 443.25 | 1.39 | 90.13 | 0.03 | 3.29 | 87.19 | 11.97 | 1.16 |
| Manus Basin | Solwara 7 | 9.95 | 11.06 | 52.63 | 26.59 | 229.26 | 12.31 | 11.89 | 0.07 | 0.59 | 29.32 | 0.89 | 435.34 | 1.77 | 88.48 | 0.03 | 3.46 | 89.66 | 12.18 | 1.01 |
| Manus Basin | Solwara 7 | 10.09 | 11.77 | 52.10 | 26.89 | 311.33 | 27.12 | 21.67 | 0.02 | 0.52 | 31.81 | 0.89 | 421.27 | 2.12 | 88.75 | 0.03 | 3.13 | 88.16 | 12.29 | 0.97 |
| Manus Basin | Solwara 7 | 10.08 | 11.20 | 53.84 | 27.15 | 178.32 | 4.70 | 4.83 | 0.05 | 0.41 | 25.74 | 0.45 | 420.91 | 1.51 | 89.59 | 0.03 | 3.16 | 90.58 | 12.68 | 0.88 |
| Manus Basin | Solwara 7 | 9.69 | 10.85 | 52.27 | 26.42 | 190.97 | 8.35 | 7.96 | 0.06 | 0.67 | 27.29 | 22.84 | 419.97 | 1.60 | 89.42 | 0.03 | 3.28 | 90.96 | 12.19 | 0.85 |
| Manus Basin | Solwara 7 | 9.74 | 10.79 | 52.54 | 26.68 | 143.85 | 2.48 | 2.44 | 0.02 | 0.25 | 25.88 | 0.63 | 426.69 | 1.39 | 88.35 | 0.03 | 2.95 | 91.19 | 12.42 | 0.79 |
| Manus Basin | Solwara 6 | 9.94 | 11.35 | 51.31 | 25.98 | 345.13 | 0.18 | 31.40 | 0.02 | 0.67 | 31.59 | 0.74 | 436.75 | 2.10 | 89.26 | 0.03 | 3.93 | 83.17 | 10.81 | 1.73 |
| Manus Basin | Solwara 6 | 9.88 | 11.44 | 52.75 | 26.76 | 300.54 | 0.26 | 26.73 | 0.04 | 1.27 | 30.57 | 0.93 | 435.47 | 1.99 | 88.00 | 0.03 | 4.57 | 86.87 | 11.13 | 1.66 |
| Manus Basin | Solwara 6 | 9.76 | 10.71 | 51.79 | 26.11 | 216.40 | 0.09 | 12.37 | 0.03 | 0.21 | 30.79 | 0.47 | 445.43 | 1.65 | 92.22 | 0.03 | 3.38 | 91.46 | 11.98 | 1.18 |
| Manus Basin | Solwara 6 | 9.92 | 11.79 | 52.57 | 26.33 | 359.53 | 0.90 | 34.02 | 0.15 | 0.48 | 32.42 | 0.75 | 442.62 | 2.17 | 89.07 | 0.03 | 3.99 | 87.48 | 10.37 | 1.79 |
| Manus Basin | Solwara 6 | 9.99 | 10.90 | 52.39 | 27.51 | 166.44 | 0.11 | 6.25 | 0.05 | 1.92 | 24.34 | 0.41 | 422.55 | 1.43 | 88.92 | 0.03 | 3.23 | 92.83 | 12.43 | 1.79 |
| Manus Basin | Solwara 6 | 10.08 | 11.08 | 53.67 | 27.17 | 215.61 | 0.15 | 12.76 | 0.04 | 1.04 | 26.52 | 0.42 | 428.51 | 1.64 | 91.18 | 0.03 | 3.65 | 88.12 | 11.67 | 1.22 |
| Manus Basin | North Su | 10.39 | 12.96 | 52.63 | 26.03 | 673.76 | 1.67 | 21.88 | 0.04 | 25.22 | 47.56 | 2.01 | 437.70 | 3.84 | 92.02 | 0.06 | 4.15 | 61.00 | 8.09 | 2.35 |
| Manus Basin | North Su | 10.21 | 11.90 | 52.84 | 25.34 | 483.66 | 2.02 | 14.94 | 0.04 | 45.68 | 40.11 | 2.02 | 429.14 | 3.07 | 91.67 | 0.06 | 3.68 | 71.41 | 9.42 | 2.26 |
| Manus Basin | North Su | 10.22 | 11.63 | 53.21 | 25.70 | 372.66 | 1.20 | 11.10 | 0.04 | 1.67 | 36.48 | 0.80 | 417.24 | 2.61 | 90.58 | 0.03 | 3.39 | 76.15 | 10.14 | 1.65 |
| Manus Basin | South Su | 9.52 | 10.80 | 53.90 | 26.69 | 195.39 | 0.28 | 2.62 | 0.07 | 22.93 | 25.08 | 1.48 | 370.88 | 1.48 | 89.68 | 0.06 | 2.86 | 88.92 | 12.24 | 1.28 |
| Manus Basin | South Su | 9.51 | 10.80 | 52.56 | 26.63 | 202.32 | 0.18 | 2.15 | 0.05 | 31.93 | 25.61 | 1.50 | 379.89 | 1.47 | 90.20 | 0.06 | 2.76 | 90.53 | 12.41 | 1.41 |
| Manus Basin | South Su | 9.48 | 10.61 | 51.22 | 24.64 | 217.31 | 0.27 | 5.66 | 0.04 | 17.40 | 26.31 | 5.17 | 406.67 | 1.60 | 84.41 | 0.04 | 2.81 | 80.86 | 10.49 | 1.72 |
| Manus Basin | South Su | 9.64 | 10.64 | 52.15 | 26.56 | 154.31 | 0.70 | 1.45 | 0.06 | 10.31 | 26.72 | 2.69 | 370.64 | 1.44 | 89.70 | 0.04 | 2.74 | 88.65 | 12.78 | 1.81 |
| Manus Basin | South Su | 10.25 | 10.82 | 54.82 | 27.85 | 150.79 | 0.27 | 1.07 | 0.07 | 24.86 | 23.95 | 2.02 | 415.77 | 1.38 | 89.19 | 0.07 | 3.13 | 88.71 | 12.10 | 0.90 |
| Manus Basin | South Su | 10.16 | 10.92 | 55.39 | 27.56 | 138.18 | 1.23 | 0.46 | 0.04 | 2.00 | 23.03 | 1.04 | 415.88 | 1.32 | 88.76 | 0.03 | 2.86 | 89.93 | 12.09 | 0.38 |
| Manus Basin | South Su | 9.69 | 10.66 | 54.27 | 27.10 | 143.31 | 0.10 | 0.40 | 0.04 | 7.00 | 24.86 | 1.13 | 371.47 | 1.35 | 91.47 | 0.04 | 2.80 | 92.36 | 12.84 | 1.03 |
| Woodlark Basin | La Scala | 9.78 | 10.48 | 52.05 | 25.43 | 209.28 | 0.08 | 6.99 | 0.05 | 1.66 | 34.71 | 0.64 | 388.90 | 1.33 | 85.40 | 0.03 | 2.17 | 84.80 | 11.68 | 0.86 |
| Woodlark Basin | La Scala | 9.60 | 10.50 | 52.89 | 26.10 | 142.21 | 0.08 | 1.34 | 0.04 | 19.73 | 26.55 | 1.08 | 377.44 | 1.27 | 85.43 | 0.05 | 2.39 | 89.72 | 12.16 | 0.88 |
| Woodlark Basin | La Scala | 9.93 | 10.70 | 54.19 | 26.62 | 114.08 | 0.09 | 0.48 | 0.04 | 2.78 | 24.94 | 1.02 | 385.01 | 1.29 | 87.31 | 0.04 | 2.31 | 93.39 | 13.13 | 0.75 |

**Supplementary Information S6**

**Boxplots for each element measured in the shells of *Shinkailepas tollmanni* encapsulated larvae.**

For each element, the number of samples (n) is indicated for all sites. When relevant, the number of values under the limit of detection (LOD) is also indicated. These represent 124 values from the 7800 of the dataset.

Several elements present different distributions between sites; this is the case for Mn, Ba and Pb, with other elements that have less obvious differences such as Mg, Fe, Zn, and Sr. Other elements have relatively similar distributions in all sites: Cr, Cu, As, Sn, Cd, and Sb. Therefore, these last elements do not appear to be promising candidates for discrimination. As five of these six elements are those presenting some values below LOD, we are able to focus on the other elements and use all 600 analysed larval shells for classification, with no values below LOD.

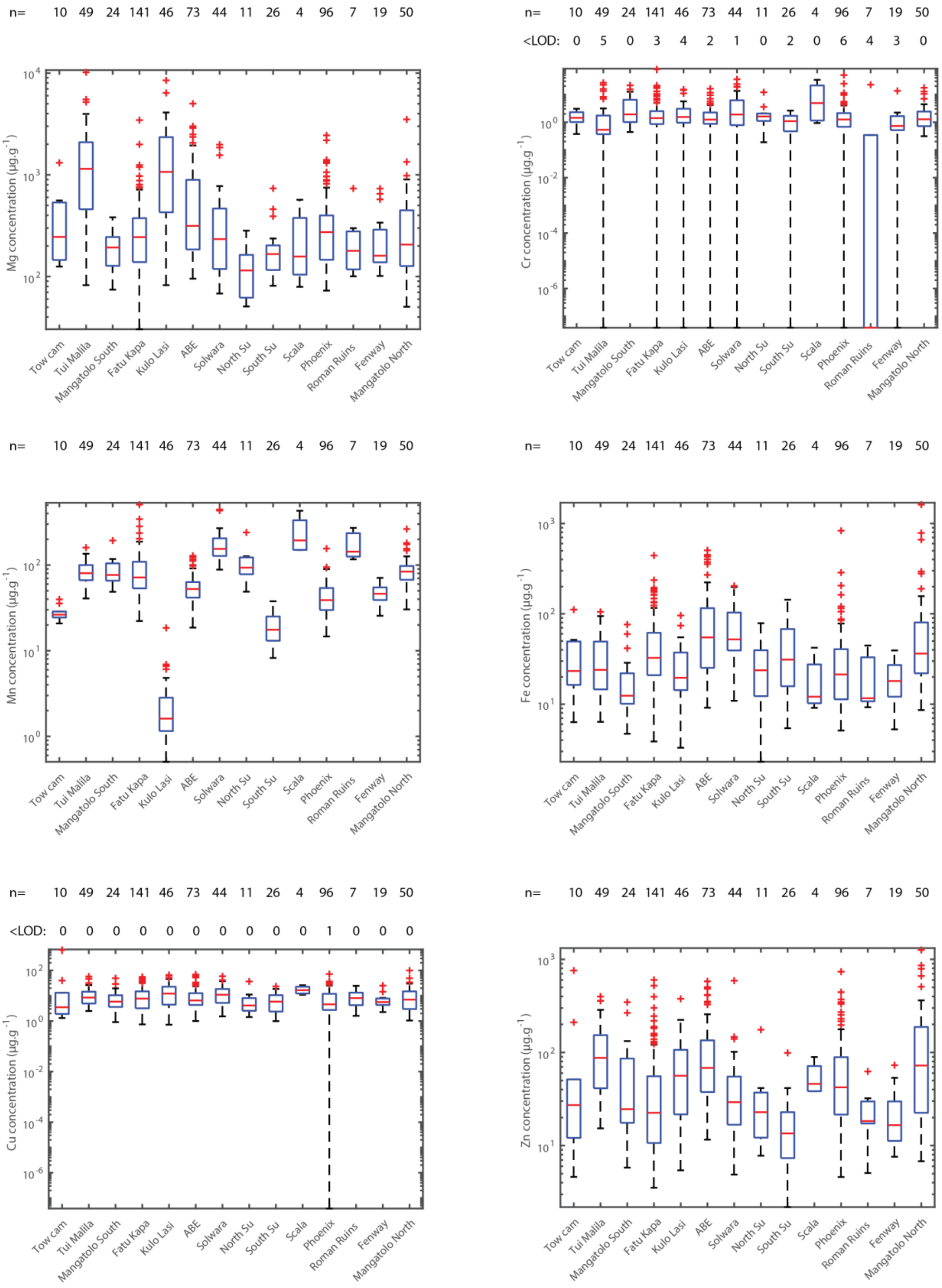


**Figure S3:** Boxplots of elemental abundance for Mg, Cr, Mn, Fe, Cu, and Zn.

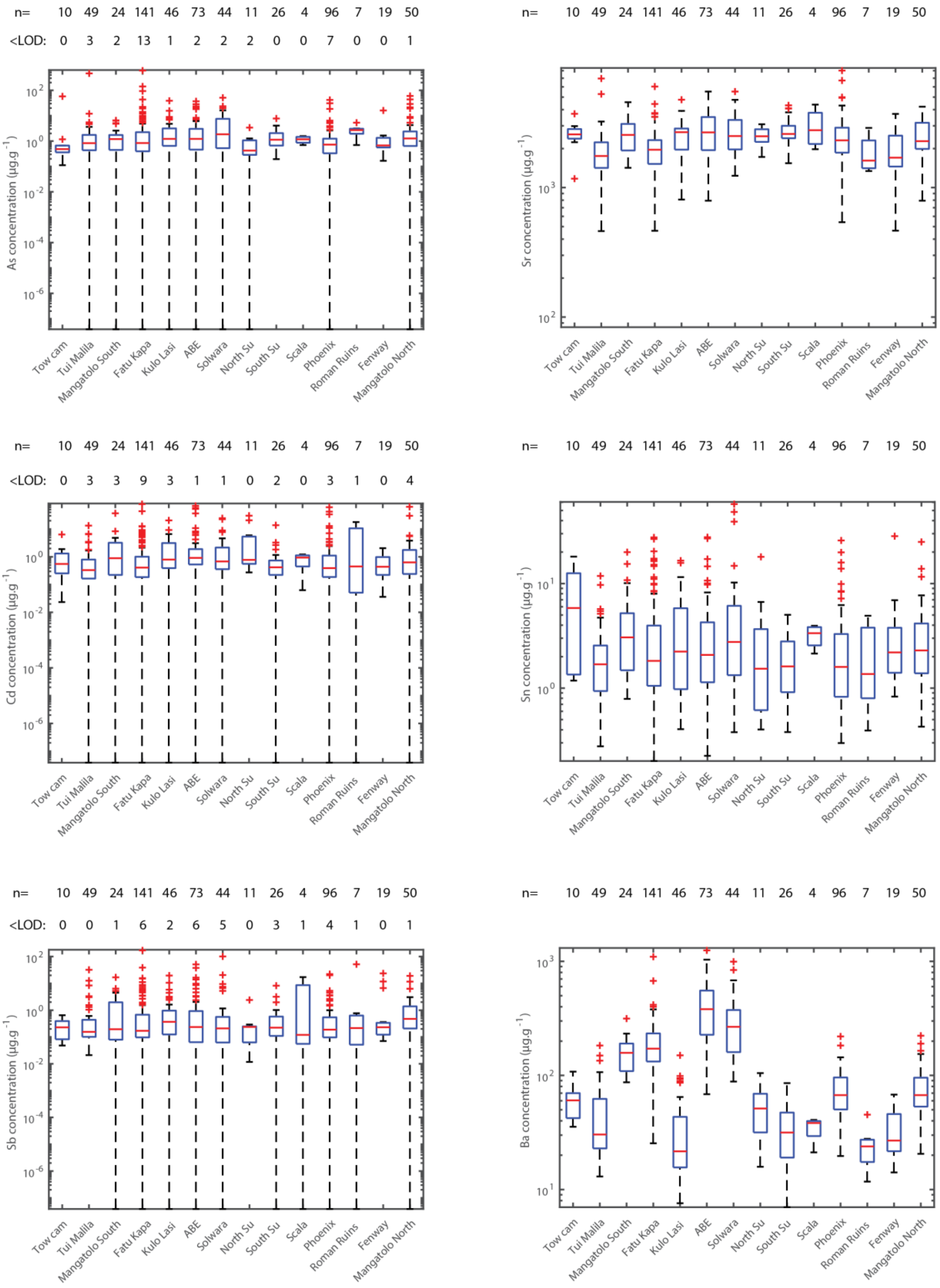


**Figure S4:** Boxplots of elemental abundance for As, Sr, Cd, Sn, Sb, and Ba.

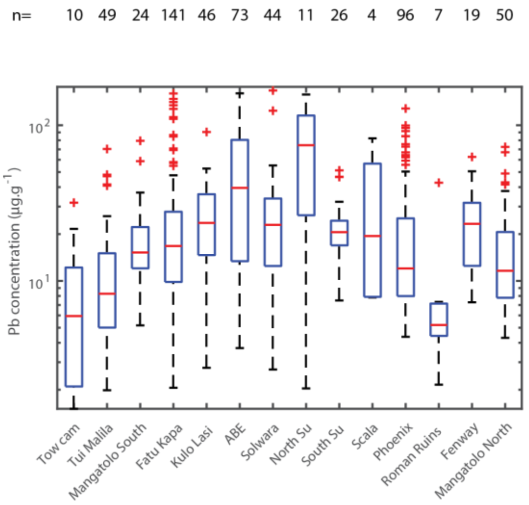


**Figure S5:** Boxplot of elemental abundance for Pb.

**Supplementary Information S7**

**Details on the larval shell elemental distributions in the Lau area**

Some collection sites were sampled several times (bioboxes) in the *Ifremeria* communities to study the variability within site. An example for Lau is given on Fig. S6. We observed no significant difference in the variability at this scale for both Tu'i Malila and ABE. The low number of specimens from Tow Cam (n=10), particularly for biobox 2 (n=2), prevents a reliable interpretation. Still, bioboxes from the same site appear generally similar, and remain different than specimens from other sites.

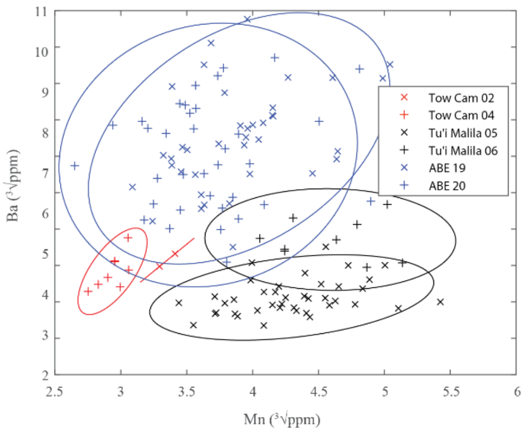


**Figure S6:** Distribution of all larval shell Mn and Ba from Lau for each site and biobox (2 bioboxes at each site).
